# *Staphylococcus aureus* Breast Implant Infection Isolates Display Recalcitrance to Antibiotic Pocket Irrigants

**DOI:** 10.1101/2022.07.18.500563

**Authors:** Jesus M. Duran Ramirez, Jana Gomez, Blake Hanson, Taha Isa, Terence Myckatyn, Jennifer N Walker

**Author notes:** Address correspondence to Jennifer N. Walker, PhD,. Jennifer N. Walker, The University of Texas Health Science Center, Houston, TX, USA. Author order was determined on the basis contribution.

## Abstract

Breast implant-associated infections (BIAIs) are a common complication following breast prostheses placement and account for ∼100,000 infections annually. The frequency, high cost of treatment, and morbidity make BIAIs a significant health burden for women. Thus, effective BIAI prevention strategies are urgently needed. This study tests the efficacy of one infection prevention strategy: the use of a triple antibiotic pocket irrigant (TAPI) against *Staphylococcus aureus*, the most common cause of BIAIs. TAPI, which consists of 50,000 U bacitracin, 1 g cefazolin, and 80 mg gentamicin diluted in 500 mL of saline, is used to irrigate the breast implant pocket during surgery. We used *in vitro* and *in vivo* assays to test the efficacy of each antibiotic in TAPI, as well as TAPI at the concentration used during surgery. We found that planktonically grown *S. aureus* BIAI isolates displayed susceptibility to gentamicin, cefazolin, and TAPI. However, TAPI treatment enhanced biofilm formation of BIAI strains. Furthermore, we compared TAPI treatment of a *S. aureus* reference strain (JE2) to a BIAI isolate (117) in a mouse BIAI model. TAPI significantly reduced infection of JE2 at 1- and 7-days post infection (dpi). In contrast, BIAI strain 117 displayed high bacterial burdens in tissues and implants, which persisted out to 14-dpi despite TAPI treatment. Lastly, we demonstrated that TAPI was effective against *P. aeruginosa* reference (PAO1) and BIAI strains *in vitro* and *in vivo*. Together, these data suggest *S. aureus* BIAI strains employ unique mechanisms to resist antibiotic prophylaxis treatment and promote chronic infection.

## Introduction

Nearly 300,000 breast prostheses are placed annually in the US for cosmetic and reconstructive purposes, of which 1%-35% become infected (1-3). These infections can cause significant patient morbidity, as they can result in pain, fluid collections, fever, tissue necrosis, and device failure (1, 3-11). Notably, most breast implant-associated infections (BIAIs) occur during reconstructive breast implant-based surgery following mastectomy due to cancer, which have infection rates as high as 35% (1-3). For women with cancer, BIAI can lead to additional complications and delay of adjuvant therapies, such as chemotherapy and radiation. Furthermore, treatment of BIAIs require the explantation of the infected prosthesis, which necessitates additional surgical procedures and administration of broad-spectrum antibiotics, increasing the risk of disseminated infection and escalating health care costs (12, 13). Thus, BIAIs are a significant health burden for women and prevention has become a priority (2, 5, 14-16).

Current evidence-based prophylaxis strategies involve the administration of preoperative intravenous antibiotics – typically first-or second-generation cephalosporins – and surgical skin scrubs at the incision site (17, 18). Yet, infection rates remain high. Thus, additional preventive strategies that involve the flushing of the surgical pocket with a triple antibiotic pocket irrigant (TAPI) have been implemented by surgeons in an effort to further reduce infection rates (17, 19). TAPIs typically consist of 50,000 U bacitracin, 1 g cefazolin, and 80 mg gentamicin diluted in 500 mL of saline and are introduced into the surgical pocket prior to breast prosthesis placement (15, 16). Recent *in vitro* studies suggest prophylactic antibiotic irrigation strategies, such as TAPI, may be efficacious against some of the most common causes of BIAI, including *Staphylococcus aureus* and *Pseudomonas aeruginosa* (4, 6, 16, 20-23). While these studies are an important first step, the majority of this work relies on minimum inhibitory concentration (MIC) assays to assess the susceptibility patterns of reference strains to TAPI or its components (2, 4, 5, 14, 16, 20, 24-34). However, reference strains may not reflect the diversity of antimicrobial resistance or virulence mechanisms carried by pathogens currently circulating in the clinic today. Furthermore, *in vitro* conditions do not always recapitulate interactions that occur during infection. Specifically, for *S. aureus*, in addition to genetic resistance, these bacteria also exhibit phenotypic drug recalcitrance (35-41). Phenotypic recalcitrance among *S. aureus* strains typically involves the incorporation of various host factors present in the blood or during wound healing, such as fibrinogen and collagen, into biofilm structures (1, 28, 42). These biofilms provide recalcitrance to antibiotic concentrations at which their planktonic form is susceptible (4, 35, 36, 40, 41, 43-49). Moreover, these host proteins are available during breast surgery, and may affect prophylactic antibiotic efficacy (24-27, 36, 43). Additionally, while TAPI prophylaxis has been associated with a reduction in the rate of capsular contracture, a latent complication thought to occur when low levels of bacteria contaminate host-formed capsules surrounding the prostheses, there is a lack of randomized, well controlled, clinical studies assessing the efficacy of TAPI at preventing overt BIAI (1, 2, 5, 9, 10, 15, 16, 50). Thus, it remains unclear how MIC assays using reference strains can be translated into effective irrigant strategies that prevent BIAIs.

In this study, we used *in vitro* and *in vivo* assays to assess the efficacy of TAPI against clinically relevant isolates of *S. aureus*, to provide insights into how the pathogen resists antibiotics to become one of the most common causes of BIAI (1, 5). We characterized two *S. aureus* BIAI isolates (117 and 158), as well as JE2, a well-studied, reference strain. Using *in vitro* MIC assays we found that all three *S. aureus* isolates displayed resistance to bacitracin, but were susceptible to cefazolin and TAPI. Notably, JE2 also displayed resistance to gentamicin, while both BIAI isolates were susceptible to the antibiotic. Surprisingly, while biofilm formed by JE2 was not affected by exposure to TAPI *in vitro*, the irrigant enhanced the biomass of the BIAI strains compared to untreated controls. Furthermore, using a mouse model, we demonstrated that the *S. aureus* BIAI strain 117 displayed increased recalcitrance to TAPI compared to JE2. Specifically, JE2 was significantly reduced by TAPI prophylaxis compared to saline treated controls at 1- and 7-days post infection (dpi), while the irrigant had no effect on the ability of the BIAI strain 117 to persist within tissues and on implants at the same time points. Notably, this phenotypic recalcitrance *in vivo* was unique to *S. aureus* clinical isolates, as when reference (PAO1) and BIAI isolates (157 and 160) of *P. aeruginosa* were assessed *in vitro* and in the mouse model, TAPI was effective at killing the strains and preventing chronic infection. Thus, these data emphasize the disparity between clinical and reference isolates when determining the efficacy of prevention strategies. Furthermore, this study highlights how *S. aureus* BIAI isolates display unique mechanisms to resist antibiotics during BIAI.

## Results

### *S. aureus* Antibiotic Susceptibility Patterns

Two *S. aureus* strains isolated from BIAIs (117 and 158) and a reference strain (JE2) (**Table 1**) were assessed for their susceptibility to gentamicin, cefazolin, bacitracin, as well as TAPI, via MIC and minimum bactericidal concentration (MBC) assays (**Figure 1 and Table S1**). Both BIAI strains 117 and 158 display susceptibility to gentamicin, while JE2 exhibits resistance as expected (51) (**Figure 1A**). All *S. aureus* strains display the same MBC of 1.0 ug/mL (**Table S1**). All *S. aureus* strains are susceptible to cefazolin, and display MBCs of 6.0, 0.5, 1.0 ug/mL for JE2, 117, and 158, respectively (**Figure 1B and Table S1**). Additionally, all *S. aureus* strains display resistance to bacitracin, and exhibit MBCs of >32 ug/mL (**Figure 1C and Table S1**). We also assessed the susceptibility of all three *S. aureus* strains to the TAPI at the concentrations used in patients. TAPI was effective against all *S. aureus* strains, as there was no bacterial growth detected (**Figure 1D and Table S1**). Lastly, the genomes of the BIAI strains 117 and 158 were sequenced, the sequence types (STs) determined, and the potential antibiotic resistance genes identified. While JE2 is a ST 8, the BIAI isolates 117 and 158 were ST 39 and ST 45, respectively (**Table S2**). Furthermore, while JE2 encodes the *mecA* gene, confirming it is a methicillin resistant *S. aureus* strain, neither BIAI isolate carried *mecA*, suggesting both are methicillin susceptible (**Table S2**). Notably, the BIAI strains only encode a few genes with known roles in resistance to tetracycline (*tet-38* and *mepA/R*), supporting the MIC data above and suggesting they harbor limited acquired antibiotic resistance mechanisms.

**Table 1.**
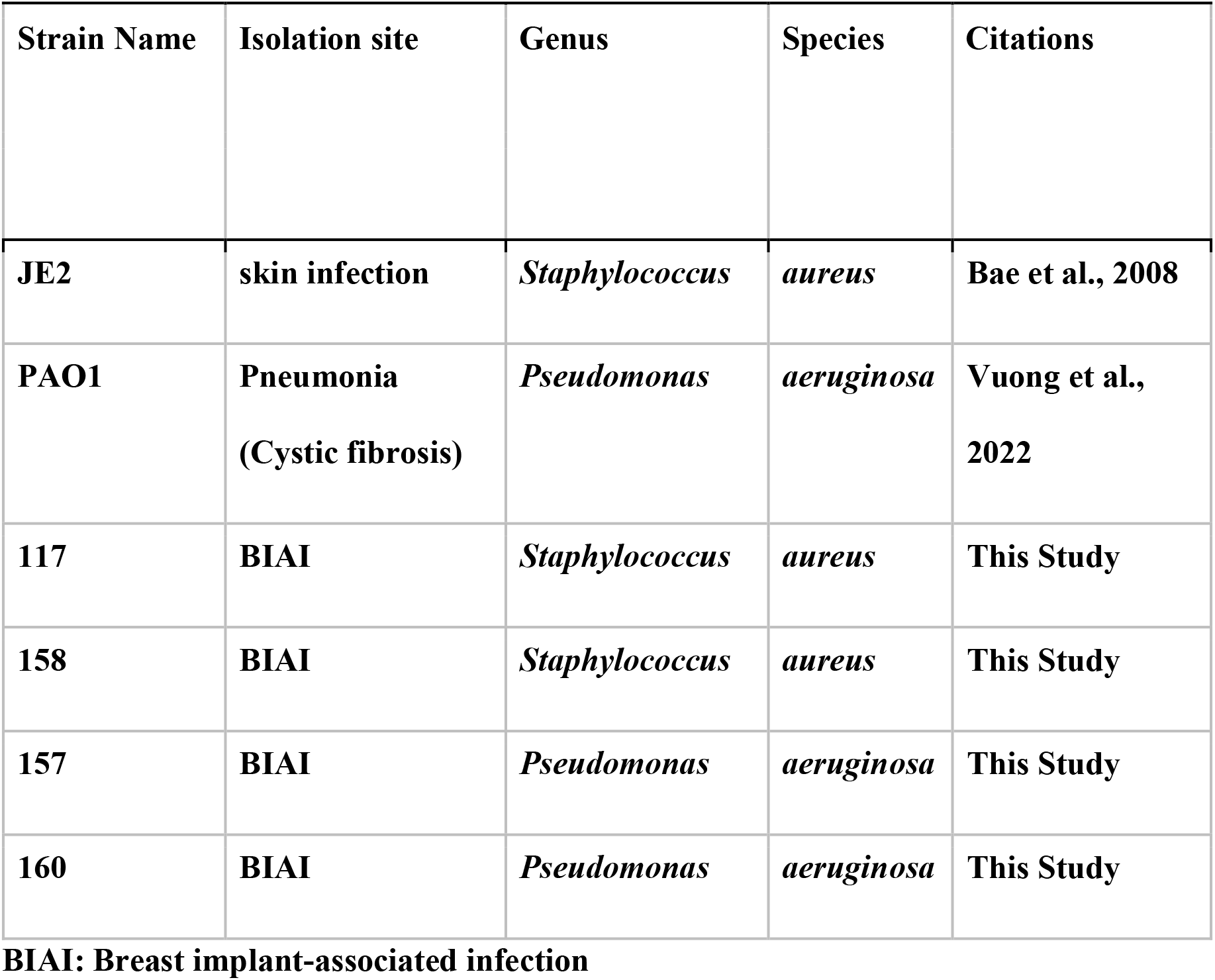
Strain information.

**Figure 1.**
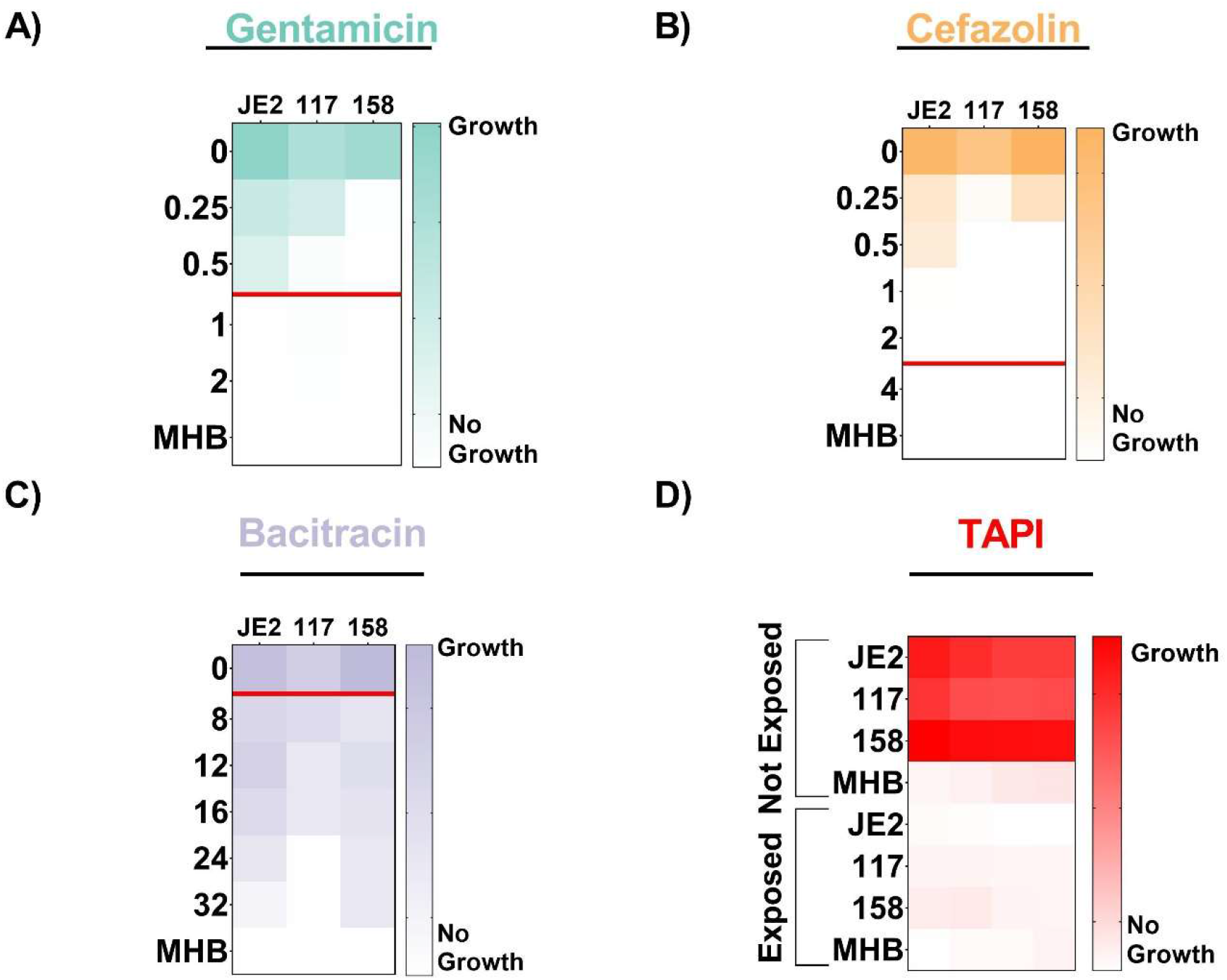
Antibiotic susceptibility patterns of *S. aureus* strains. Strains JE2, 117, and 158 were exposed to increasing concentrations of gentamicin (**A**), cefazolin (**B**), or bacitracin (**C**). **A**) JE2 displays resistance to gentamicin, with a MIC of 1.0 ug/mL, while 117 and 158 are susceptible, as they exhibit MICs of 0.5 and 0.25 ug/mL, respectively. **B**) JE2, 117, and 158 are all susceptible to cefazolin, with MICs of 1.0, 0.5, and 0.5 ug/mL, respectively. **C**) JE2, 117, and 158 are all resistant to bacitracin, as they exhibit MICs of >32, 24, and >32 ug/mL, respectively. The red line represents the MIC breakpoint for each antibiotic for *S. aureus*. A MIC value at or above the MIC breakpoint classifies the pathogen as resistant to the antibiotic. **D**) Susceptibility of *S. aureus* strains to TAPI was determined based on an increase in optical density, indicating growth of strains, when exposed to TAPI compared to those not exposed to TAPI. JE2, 117, and 158 display susceptibility to TAPI.

### *S. aureus* Biofilms Affect Antibiotic Efficacy

All three *S. aureus* strains were assessed for biofilm formation and were able to form biofilm under standard *in vitro* conditions (**Figure 2**). Additionally, biofilms were exposed to increasing concentrations of gentamicin and cefazolin, the antibiotics at which the BIAI isolates were susceptible during planktonic growth, including the MIC, twice the MIC, and four times the MIC. There was no significant difference in biomass among any of the strains when increasing concentrations of antibiotics were added to preformed biofilms (**Figure 2**). However, TAPI significantly reduced the biomass of JE2 and the BIAI strain 158 (**Figure 2A and C**). In contrast, the biomass of the BIAI strain 117 was not affected by TAPI (**Figure 2B**). Additionally, biofilms were formed in the presence of human plasma to more closely mimic infection-like conditions, as previously described (29). Human plasma significantly enhanced biofilms formed by all *S. aureus* strains compared to those grown in media alone (**Figure 2 and 3**). Additionally, when these biofilms were exposed to increasing concentrations of gentamicin and cefazolin there was no significant difference in biomass formed by any of the strains compared to unexposed controls, similarly to biofilms formed in media alone (**Figure 3A, B, and C**). However, in contrast to biofilms formed in media alone, TAPI had no effect on the biomass formed by JE2 under these conditions (**Figure 3A**). Notably, TAPI significantly increased the biomass formed by both of the BIAI isolates compared to control, unexposed biofilms under these conditions (**Figure 3B and C**).

**Figure 2.**
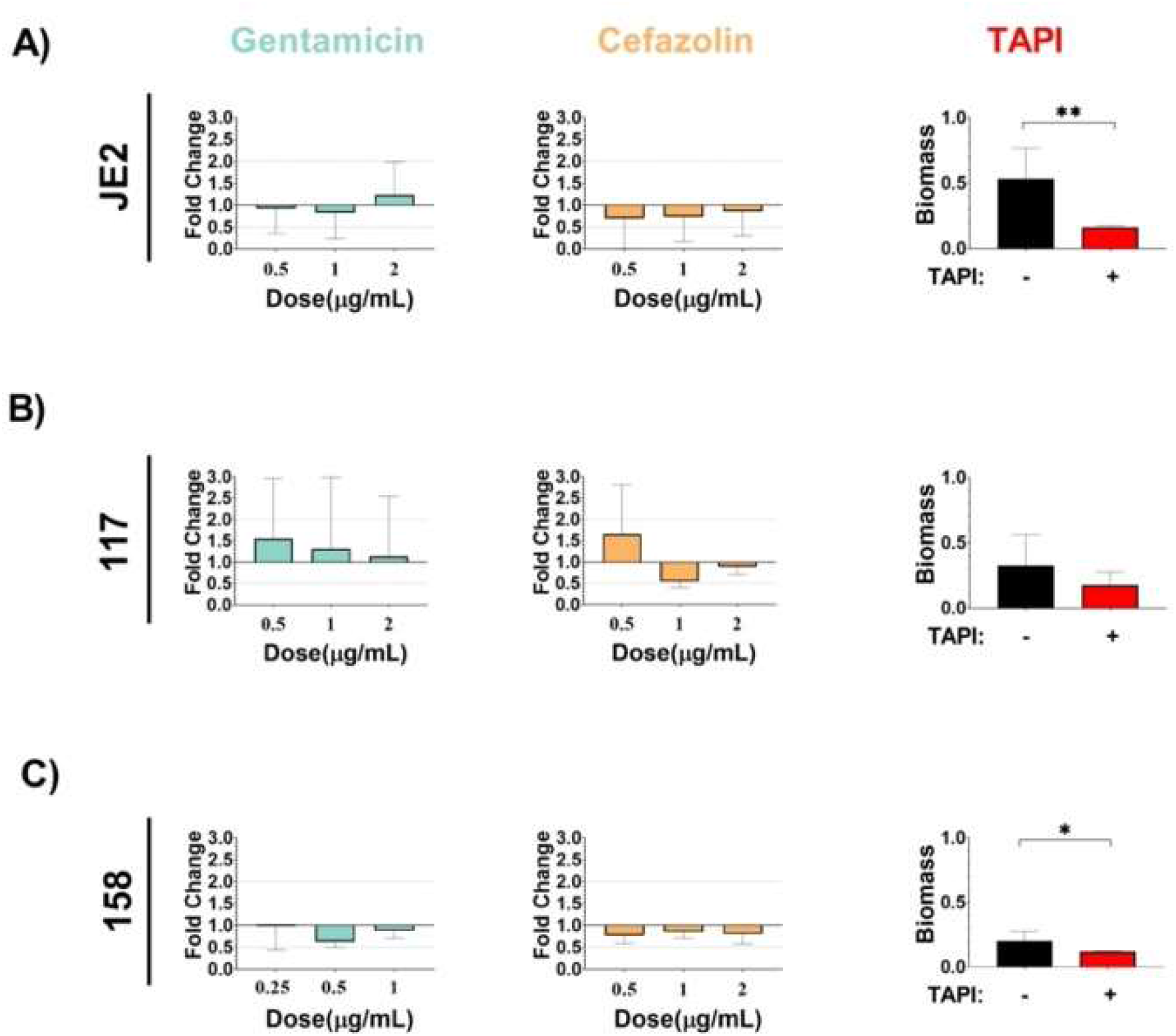
*S. aureus* biofilms formed under *in vitro* conditions display recalcitrance to TAPI antibiotics. JE2 (**A**), 117 (**B**), and 158 (**C**) were allowed to form biofilm and were then exposed to increasing concentration of antibiotics. **A**) JE2 biofilm was not significantly affected following exposure to increasing concentrations of gentamicin or cefazolin. TAPI, however, significantly reduced the amount of biomass formed by JE2. **B**) 117 biofilm was not significantly affected following exposure to increasing concentrations of gentamicin or cefazolin or to TAPI. **C**) 158 biofilm was not significantly affected following exposure to increasing concentrations of gentamicin or cefazolin. However, TAPI significantly reduced the amount of biomass formed by 158.

**Figure 3.**
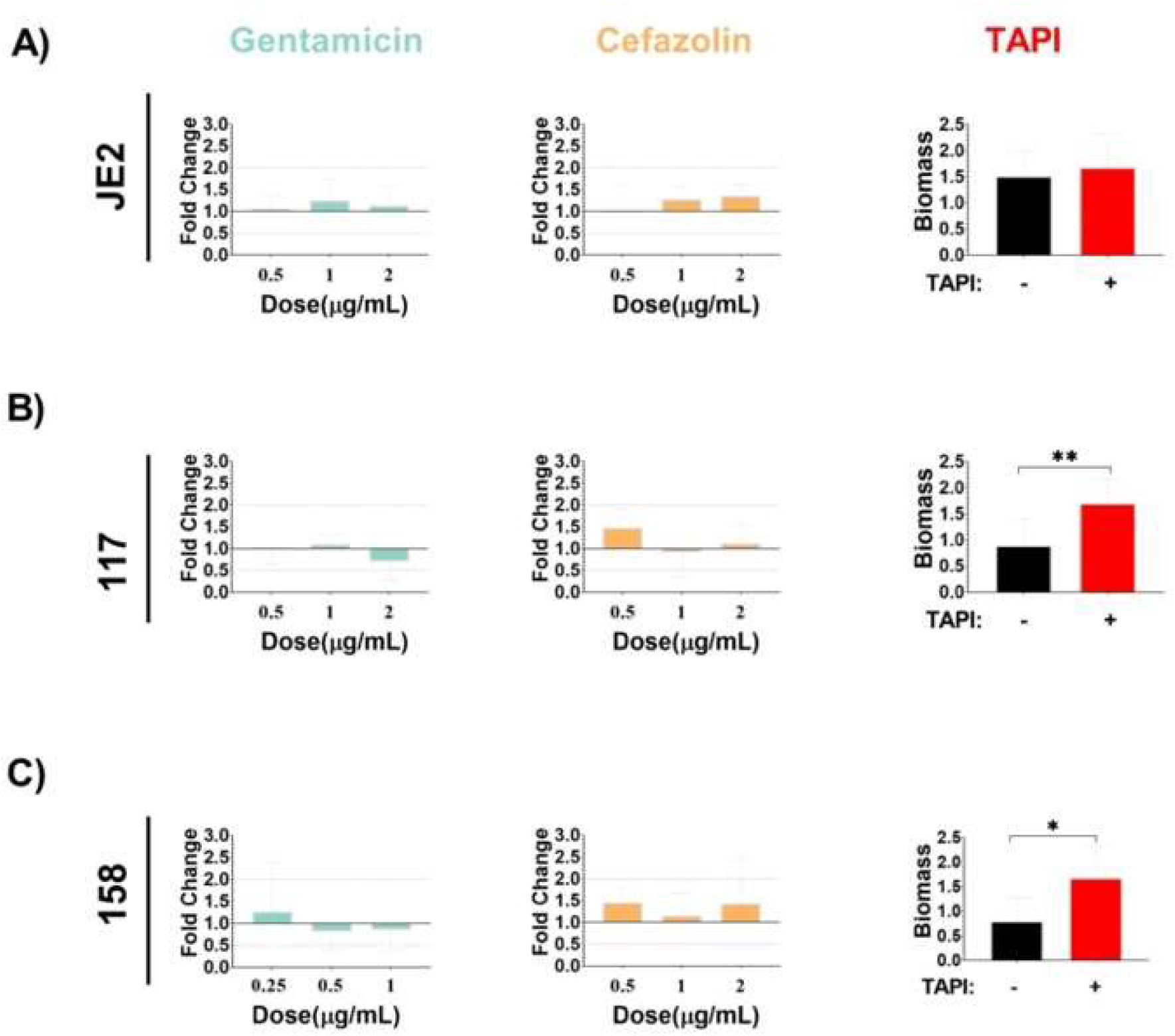
*S. aureus* biofilms formed under *in vitro* conditions that mimic the host environment display recalcitrance to TAPI antibiotics. JE2 (**A**), 117 (**B**), and 158 (**C**) were allowed to form biofilm in the presence of human plasma and were then exposed to increasing concentrations of antibiotics. **A**) JE2 biofilm was not significantly affected following exposure to increasing concentrations of gentamicin or cefazolin or to TAPI. **B**) 117 biofilm and **C**) 158 biofilm were not significantly affected following exposure to increasing concentrations of gentamicin or cefazolin, but were significantly increased when exposed to TAPI.

### *S. aureus* Displays Recalcitrance to TAPI in a Mouse BIAI Model

One representative *S. aureus* BIAI isolate (117), as well as the reference strain (JE2), were selected and the efficacy of TAPI against these strains was assessed in a mouse model of BIAI (**Figure 4**). For saline treated control mice, high JE2 colony forming units (CFUs) were recovered from implants (3.8×10^5^ CFUs) and the corresponding tissue samples (3.4×10^9^ CFUs) harvested at 1-dpi (**Figure 4A and B**). This infection persisted at 7-dpi, as similar CFUs were recovered from implants (5.3×10^5^ CFUs) and tissues (3.3×10^8^ CFUs) (**Figure 4A and B**). However, at 14-dpi only one of the recovered implants and about half of the tissues had detectable JE2 CFUs (**Figure 4A and B**). Compared to control mice, TAPI significantly reduced the JE2 CFUs in the tissue at both 1- and 7-dpi, with 6.7×10^8^ and 1.9×10^7^ CFUs recovered, respectively (**Figure 4B**). At 14-dpi, about half of the TAPI treated mice cleared JE2 from the tissue, similar to control mice. However, for implants, TAPI only significantly reduced the JE2 CFUs at 1-dpi (**Figure 4A**). There were no differences detected in JE2 CFUs recovered from implants of TAPI treated compared to control mice at 7-or 14-dpi.

**Figure 4:**
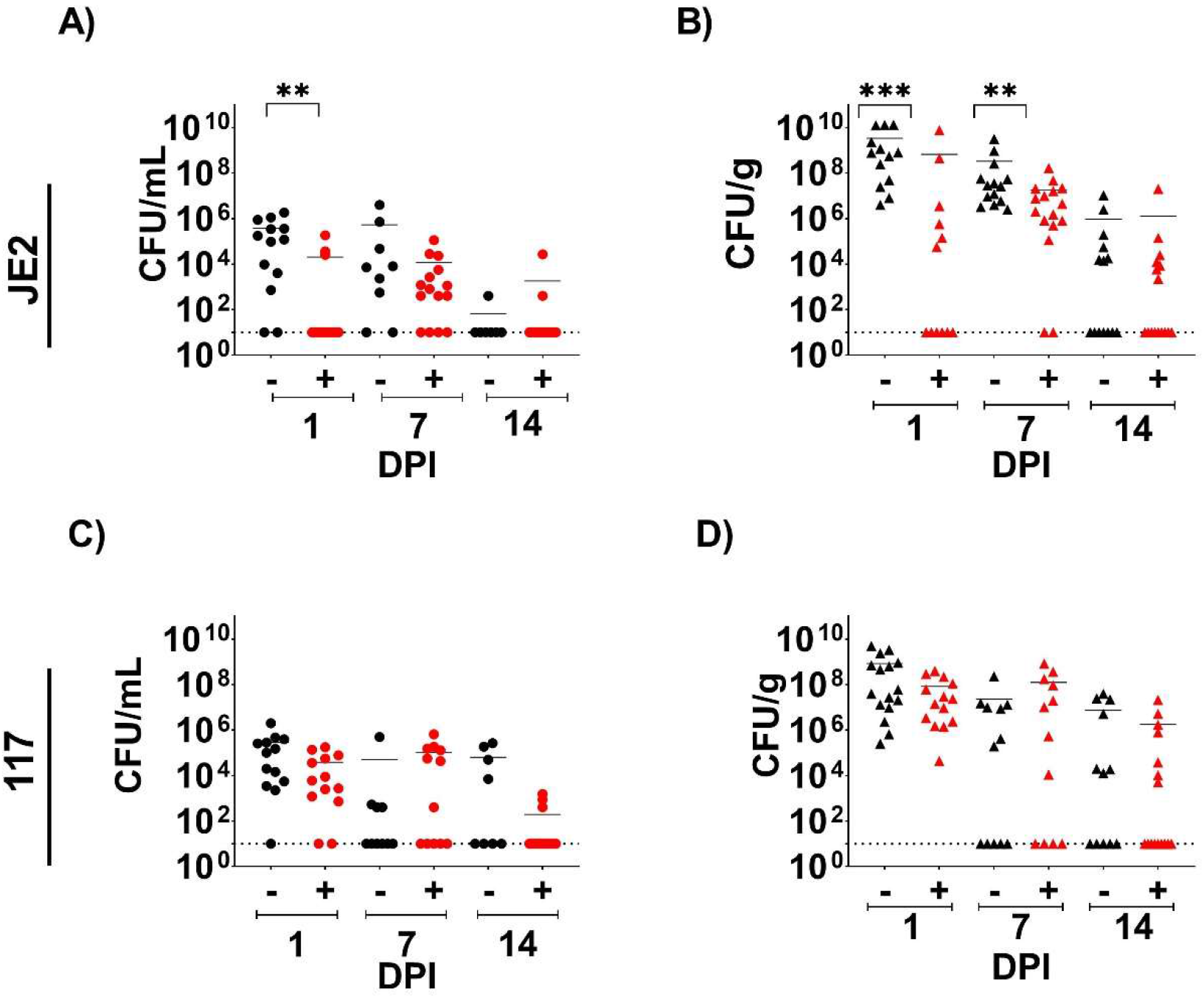
Mouse model of *S. aureus* BIAIs. **A**) Implants recovered from saline treated mice displayed high colony forming units (CFUs) of JE2 at 1 day post infection (dpi), which persisted at 7-dpi. However, mice were able to control JE2 infection by 14-dpi. TAPI significantly reduced JE2 CFUs on implants compared to the saline treated control mice at 1-dpi. However, by 7-dpi there was no difference in CFUs recovered from TAPI or saline treated mice. At 14-dpi, most mice cleared JE2 regardless of treatment. **B**) Tissue from saline treated mice displayed high CFUs of JE2 at 1- and 7-dpi. However, half the mice were able to control the infection by 14-dpi. TAPI significantly reduced CFUs in the tissue compared to the saline treated mice at 1- and 7-dpi. Again, about half the mice cleared the infection with JE2 regardless of treatment by 14-dpi. **C**) Implants recovered from saline treated mice displayed high CFUs of 117 at 1-dpi, which persisted at both 7- and 14-dpi. There was no difference in CFUs recovered from implants of TAPI or saline treated mice at any time point. **D**) Tissue from saline treated mice displayed high CFUs of 117 at 1-dpi, which persisted at both 7- and 14-dpi. There was no difference in CFUs recovered from tissues of TAPI or saline treated mice at any time point. Red represents TAPI treated mice, while black denotes saline treated mice. Circles represent the CFUs recovered from implants. Triangles denote the CFUs retrieved from tissue near the implant. The Mann Whitney U test was used to determine statistical significances, with **=p<0.01, and ***=p<0.0005.

For the BIAI strain 117, high CFUs were recovered from implants (2.8×10^5^ CFUs) and the corresponding tissue samples (8.5×10^8^ CFUs) harvested at 1-dpi from control mice (**Figure 4C and D**). Notably, this infection persisted at 7- and 14-dpi, as similar CFUs were recovered from implants (5.0×10^4^ CFUs and 6.3×10^4^ CFUs, respectively) and tissues (2.3×10^7^ CFUs and 7.6×10^6^ CFUs, respectively) (**Figure 4C and D)**. Importantly, TAPI was ineffective at reducing the CFUs within either the implants or tissues of the mice infected with the BIAI strain 117 at any time point tested **(Figure 4C and D)**. Thus, while TAPI treatment is effective against JE2, which is cleared by the mice over time, the BIAI 117 strain resists the irrigant resulting in chronic infection.

### *P. aeruginosa* Antibiotic Susceptibility Patterns

To determine whether TAPI recalcitrance was unique to *S. aureus* BIAI isolates or whether other species causing these infections also display a similar phenotype, TAPI efficacy was assessed against *P. aeruginosa* strains (**Table 1**). *P. aeruginosa* BIAI isolates (157 and 160) and a reference strain (PAO1) were assessed for their susceptibility to gentamicin, cefazolin, bacitracin, and TAPI, via MIC and MBC assays (**Figure 5 and Table S1)**. All of the *P. aeruginosa* strains display susceptibility to gentamicin, and exhibit MBCs of 6 ug/mL, 4 ug/mL, and 6 ug/mL for PAO1, 157, and 160, respectively (**Figure 5A and Table S1**). However, all *P. aeruginosa* strains are resistant to cefazolin and bacitracin, and display MBCs of >32 ug/mL for cefazolin and >1400 ug/mL for bacitracin for all three isolates (**Figure 5B, C, and Table S1**). For all *P. aeruginosa* strains exposed to TAPI, no bacterial growth was detected, indicating the bacteria are susceptible to this combination of antibiotics *in vitro* (**Figure 5D and Table S1**). Lastly, the genomes of the BIAI strains 157 and 160 were sequenced, the STs determined, and the potential antibiotic resistance genes identified. While PAO1 is a ST 549, both BIAI strains were ST 633 (**Table S2**). Furthermore, both BIAI strains encode genes with known roles in resistance to cephalosporins (OXA-486), supporting the MIC data for cefazolin above.

**Figure 5.**
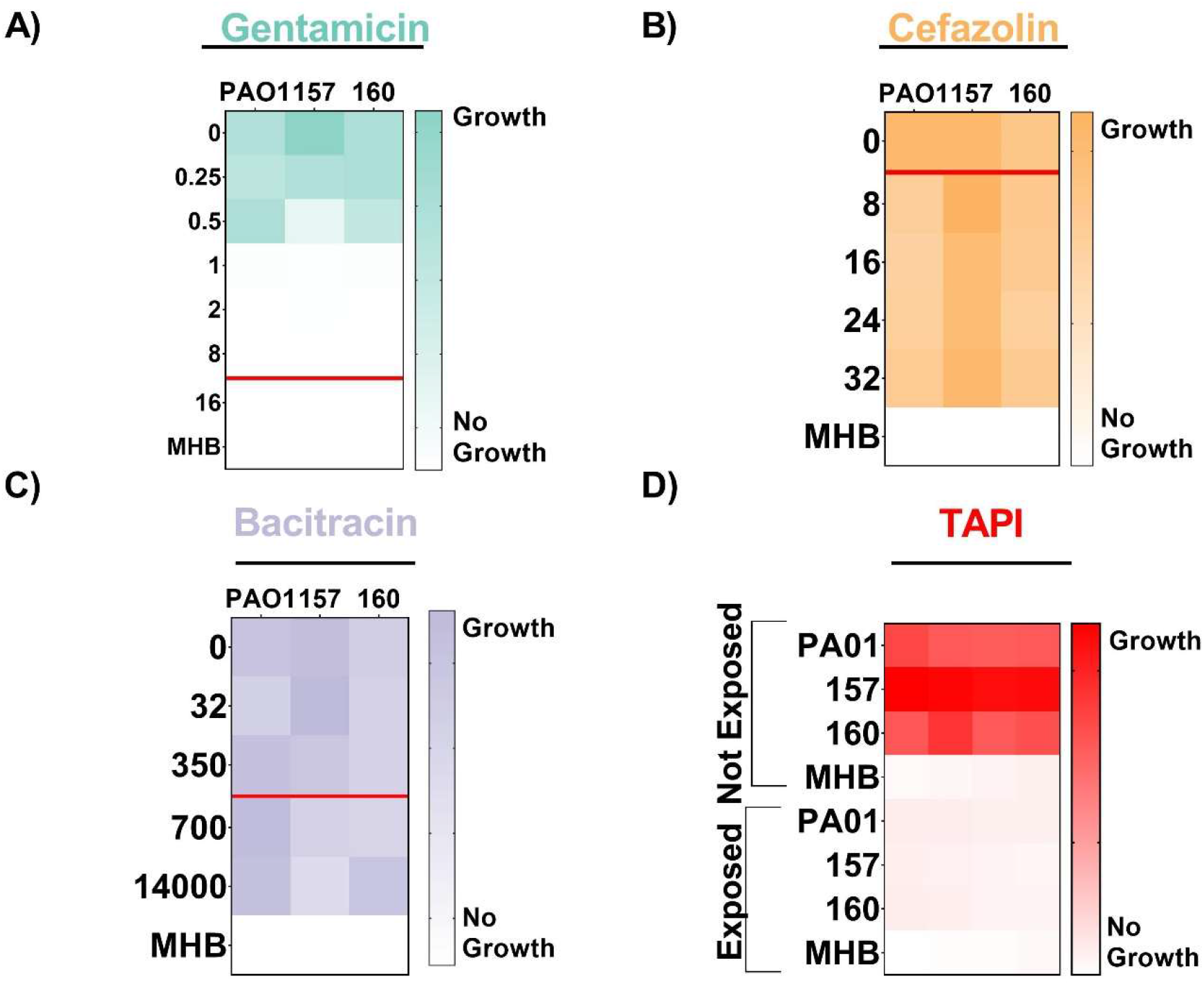
Antibiotic susceptibility patterns of *P. aeruginosa* strains. PAO1, 157, and 160 strains were exposed to increasing concentrations of gentamicin (**A**), cefazolin (**B**), or (**C**) bacitracin. **A**) PAO1, 157, and 160 all displayed minimum inhibitory concentrations (MICs) of 1 ug/mL, indicating they are susceptible to gentamicin. **B**) PAO1, 157, and 160 all displayed MICs of >32 ug/mL, indicating they are all resistant to cefazolin. **C**) PAO1, 157, and 160 all displayed MICs of >1400 ug/mL, indicating they are all resistant to bacitracin. The red line represents the MIC breakpoint for each antibiotic for *P. aeruginosa*. A MIC value at or above the MIC breakpoint classifies the pathogen as resistant to the antibiotic. **D**) Susceptibility of *P. aeruginosa* strains to TAPI was determined based on an increase in optical density, indicating growth of strains, when exposed to TAPI compared to those not exposed. PAO1, 157, and 160 are susceptible to TAPI.

### TAPI is Effective Against Communities Formed by *P. aeruginosa*

The three *P. aeruginosa* strains were assessed for biofilm and aggregate formation using previously published conditions (4, 35, 43). The *P. aeruginosa* strains were able to form biofilms and aggregates (**Figure 6**). Additionally, biofilms and aggregates were exposed to increasing concentrations of gentamicin, the antibiotic which the BIAI isolates were susceptible to during planktonic growth, including the MIC, twice the MIC, and four times the MIC. Biofilm formed by the historical strain PAO1 was significantly reduced in the presence of 2 ug/mL and 4ug/mL of gentamicin (**Figure 6A**). Additionally, aggregates were also significantly reduced in the presence of 2 ug/mL of gentamicin (**Figure 6B**). Biofilm formed by the BIAI strain 157 was also significantly reduced at 4 ug/mL of gentamicin (**Figure 6C**). However, the aggregates were unaffected by any concentration of gentamicin tested (**Figure 6D**). Lastly, while the biofilm formed by the BIAI strain 160 was recalcitrant to all the concentrations of gentamicin tested (**Figure 6E**), the aggregates were significantly reduced at 4 ug/mL (**Figure 6F**). Notably, TAPI significantly reduced biofilm formation of PAO1, 157, and 160, indicating that the antibiotics combined at the concentration used in TAPI were effective at killing *P. aeruginosa* (**Figure 6A, C, and E**).

**Figure 6:**
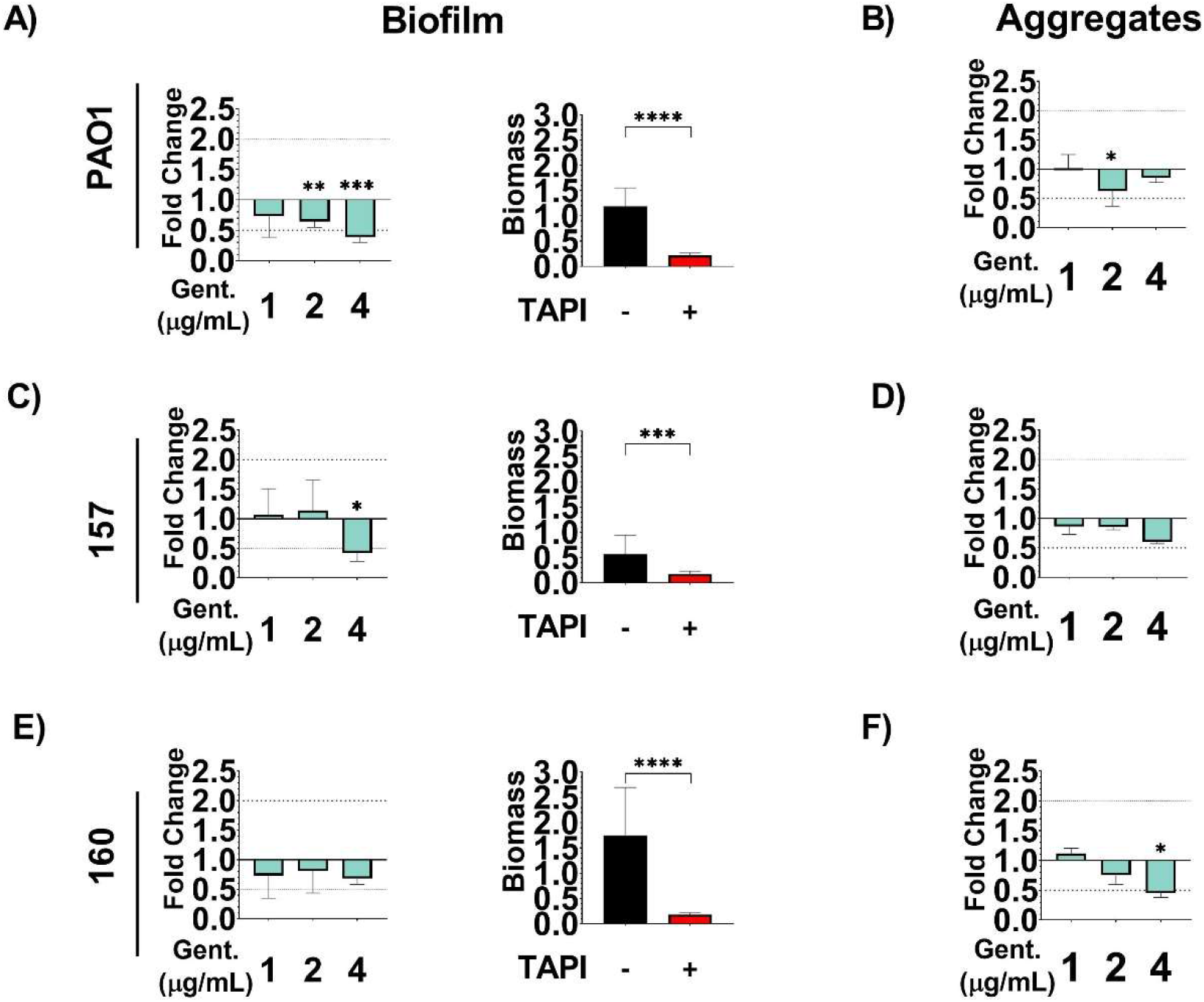
Communities formed by *P. aeruginosa* are susceptible to TAPI antibiotics. *P. aeruginosa* strains were allowed to form biofilms and aggregates and were then exposed to increasing concentrations of gentamicin, the antibiotic at which the planktonic bacteria were susceptible. **A**) 2 and 4 ug/mL (twice and four times the MIC respectively) of gentamicin significantly reduced the biomass of PAO1 biofilms. TAPI also significantly reduced PAO1 biofilm. **B**) 2 ug/mL (twice the MIC) of gentamicin significantly reduced PAO1 aggregate biomass. **C**) 4 ug/mL (four times the MIC) of gentamicin significantly reduced the biomass of the biofilm form by the BIAI strain 157. TAPI also significantly reduced the biofilm biomass. **D**) No concentration of gentamicin tested affected the biomass of aggregates formed by the BIAI strain 157. **E**) No concentration of gentamicin tested affected the biofilm formed by the BIAI strain 160. However, TAPI significantly reduced the biofilm biomass. **F**) 4 ug/mL (Four times the MIC) of gentamicin significantly reduced the biomass of the aggregates formed by the BIAI strain.

### TAPI is Effective against *P. aeruginosa* in a Mouse Model of BIAI

To investigate the efficacy of TAPI against *P. aeruginosa*, the reference strain (PAO1) and one BIAI isolate (157) were assessed in the mouse BIAI model. For control mice, high PAO1 CFUs were recovered from implants and the corresponding tissue samples at 1- and 7-dpi (**Figure 7A and B**). However, at 14-dpi control mice largely eliminated PAO1 from implants and tissues. Notably, TAPI significantly reduced the PAO1 CFUs recovered from implants and tissues at 1-dpi (**Figure 7A and B**). At 7-dpi, TAPI also significantly reduced PAO1 CFUs recovered from implants compared to control mice (**Figure 7A**). In contrast, however, PAO1 CFUs recovered from tissues of TAPI treated mice were similar to those from control mice at the same time point (**Figure 7B**). By 14-dpi, similar CFUs were recovered from both implants and tissues of TAPI treated mice compared to control mice (**Figure 7A and B**). For the *P. aeruginosa* BIAI strain 157, 1.0×10^3^ and 8.1×10^5^ CFUs were recovered from implants and the corresponding tissue samples of control mice, respectively, at 1-dpi (**Figure 7C and D**). At 7-dpi, only one implant had detectable CFUs, while more the tissue samples had detectable CFUs at the same time point (**Figure 7C and D**). At 14-dpi, only three implants and five of the corresponding tissue samples had detectable CFUs (**Figure 7C and D**). Notably, TAPI was effective at preventing infection with the BIAI strain 157, as no bacteria were recovered from implants or tissue from 1-or 14-dpi **(Figure 7C and D)**. However, at 7-dpi low CFUs of the BIAI strain 157 were recovered from only a few implants and tissue samples of TAPI treated mice **(Figure 7C and D)**. Interestingly, there was no difference between CFUs recovered from tissues of the control mice compared to the TAPI treated mice infected with either PAO1 or the BIAI strain 157 at 7-dpi (**Figure 8B and D**). Importantly, TAPI significantly reduced PAO1 and the BIAI strain 157 at early time points, with the BIAI strain 157 displaying increased susceptibility to TAPI, as the mice were successful at eliminating the infection by 14-dpi.

**Figure 7:**
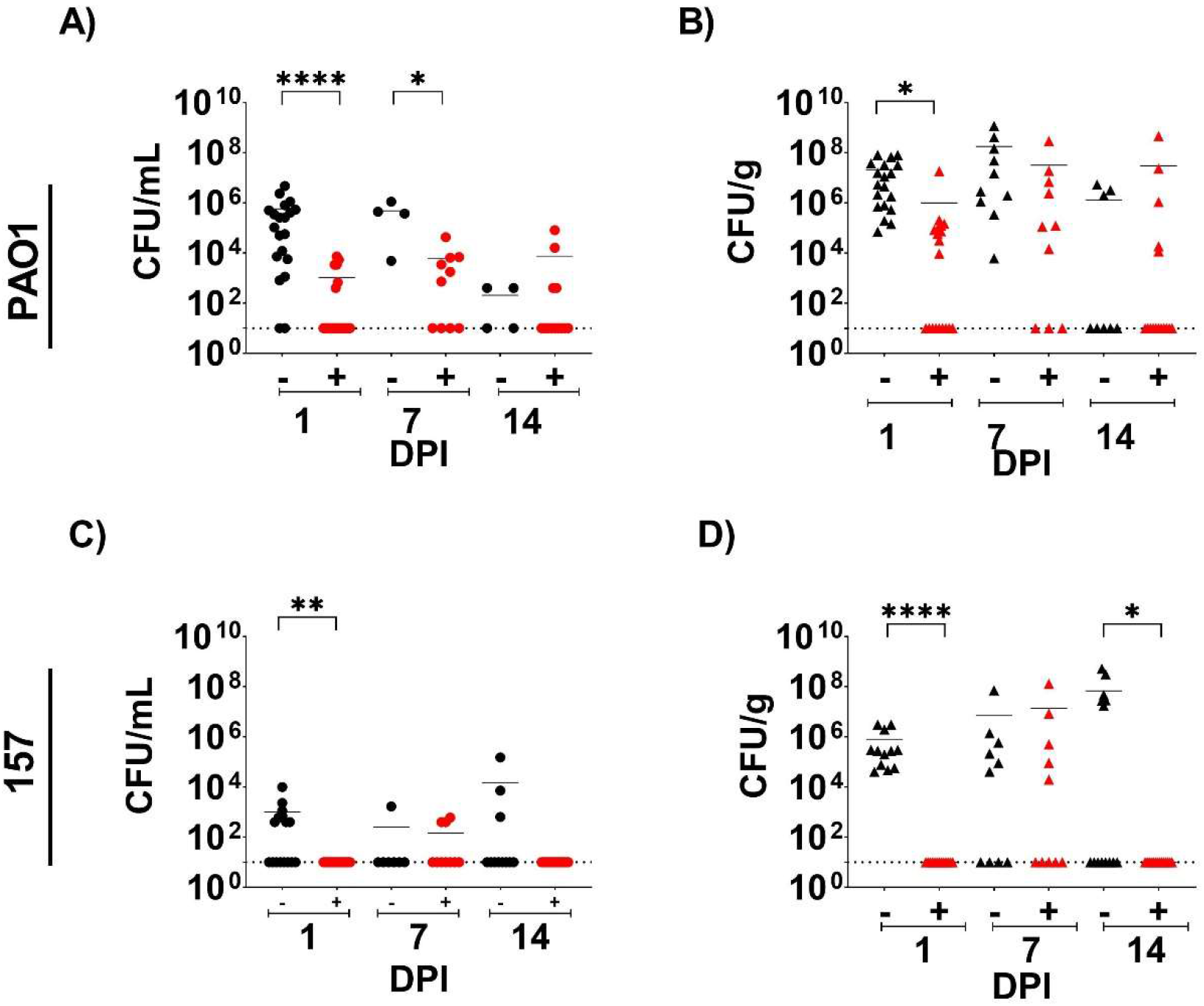
*P. aeruginosa* mouse BIAI model. **A**) Implants recovered from saline treated, control mice displayed high colony forming units (CFUs) of PAO1 at 1 day post infection (dpi), which persisted at 7-dpi. However, mice were able to eliminate PAO1 by 14-dpi. TAPI significantly reduced CFUs on implants compared to control mice at 1- and 7-dpi. At 14-dpi most mice eliminated PAO1 regardless of treatment. **B**) Tissue from control mice displayed high CFUs of PAO1 at 1-dpi, which persisted at 7-dpi. At 14-dpi, most mice eliminated PAO1 from the tissue. TAPI significantly reduced PAO1 CFUs in the tissue compared to control mice at 1-dpi. However, at 7-dpi similar CFUs were recovered from the tissue of TAPI treated compared to control mice. At 14-dpi, most TAPI treated mice eliminated PAO1 from the tissue, which was similar to control mice. **C**) CFUs of BIAI strain 157 could be detected on implants recovered from control mice at 1-dpi. However, at 7- and 14-dpi, most control mice eliminated 157 from the implants. TAPI significantly reduced the 157 CFUs compared to control mice, as no bacteria were recovered from implants of TAPI treated mice at 1- and 14-dpi, while only a few TAPI treated mice had detectable CFUs on implants at 7-dpi. **D**) Tissue from control mice displayed high CFUs of BIAI strain 157 at 1- and 7-dpi. However, more than half of the control mice had no detectable 157 CFUs at 14-dpi. TAPI significantly reduced CFUs in the tissue compared to the control mice at 1- and 14-dpi. While there was no difference in CFUs between TAPI and control mice at 7-dpi, few mice had detectable CFUs of 157 in the tissue. Red represents TAPI treated mice, while black denotes control mice. Circles signify the CFUs on implants. Triangles denote the CFUs in tissue samples near the implant. Statistical significances were determined using the Mann Whitney U test, with *=p<0.05, **=p<0.01, and ****=p<0.0001.

## Discussion

Up to a third of all breast prostheses placed annually become infected, despite sterile surgical techniques and the use of infection prevention strategies, such as surgical skin scrubs, pre- and post-operative antibiotic administration, and TAPIs (1, 2, 4-7). Furthermore, BIAIs are extremely difficult to treat, as they result in recalcitrant biofilm-associated infections that resist antibiotic therapies and require explantation of the infected prosthesis for complete resolution (1, 2, 5, 7). Thus, the implementation of evidence-based strategies that effectively prevent BIAIs have become a priority. This study focuses on assessing the efficacy of a commonly used prevention method, TAPI, against some of the most common causes of BIAIs, *S. aureus* and *P. aeruginosa*. We used recently isolated, clinically relevant strains to gain insights into the genomic and phenotypic antibiotic resistance mechanisms of currently circulating isolates causing BIAIs.

Antibiotic susceptibility testing revealed that the *S. aureus* BIAI isolates were susceptible to two of the antibiotics that make up TAPI (gentamicin and cefazolin), but resistant to the third (bacitracin). Additionally, while the *P. aeruginosa* BIAI strains were similarly resistant to bacitracin, these isolates also exhibited resistance to a second antibiotic in TAPI, cefazolin. Using whole genome sequencing (WGS) and bioinformatics analyses we found that the *S. aureus* BIAI isolates carried only a few acquired resistance genes, which mostly provide resistance to tetracycline, an antibiotic not used in TAPI (53). In contrast, the reference *S. aureus* strain JE2 encodes the *mecA* and *lmr* genes, which provide resistance to *β*-lactam and aminoglycoside antibiotics, respectively (54-56). Interestingly, all three *P. aeruginosa* strains encoded similar genes that provide resistance to cephalosporins, including *oxa*-50 for PAO1 and *oxa-486* for both BIAI strains (57-59). Thus, the genomic analyses support our MIC data. Importantly, while all the *S. aureus* and *P. aeruginosa* strains assessed exhibited resistance to at least one antibiotic, combining the drugs at the concentrations used in TAPI was effective at killing all the strains grown under planktonic conditions, which supports previous *in vitro* work (4, 14). Since the current antibiotic concentrations in TAPI are extremely high – 160, 500, and 208 times the MIC of gentamicin, cefazolin, and bacitracin, respectively, for *S. aureus*; and 10, 250, and 2 times the MIC for gentamicin, cefazolin, and bacitracin, respectively, for *P. aeruginosa –* it’s not surprising that TAPI is effective *in vitro*. However, administering an antibiotic (even at these exceedingly high concentrations) to treat infections caused by strains with known resistance to that antibiotic demonstrates a lack of antibiotic stewardship. Furthermore, while this study only tests a limited number of isolates, the fact that even the reference *S. aureus* and *P. aeruginosa* strains display resistance to bacitracin, calls into question why this antibiotic is included in TAPI. Specifically, antimicrobial stewardship dictates that if an infecting pathogen is resistant to an antibiotic or if there is a high incidence of resistance among certain patient populations, that specific antibiotic should not be used to treat that infection as treatment failure is likely to occur (60-62).

By assessing TAPI efficacy *in vivo*, our results provide support to the importance of antibiotic stewardship guidelines. Specifically, by selecting representative *P. aeruginosa* and *S. aureus* BIAI isolates and comparing the infection phenotypes to reference strains, we demonstrated that a surprising number of these bacteria could persist despite TAPI treatment. While, the *P. aeruginosa* BIAI strain 157 was the most susceptible to TAPI, and resulted in the bacteria being eliminated from implants and tissues, TAPI only significantly reduced the CFUs of the reference strains – JE2 and PAO1 – in the samples at early time points. Furthermore, almost all mice infected with either reference strain maintained high CFUs over a 7-d time course regardless of treatment. However, by 14-dpi, about half the mice began to control the infection, regardless of treatment. These results correspond with a previous study using a mouse model of skin infection that showed a different historical *S. aureus* strain followed a similar trend of spontaneously eliminating the bacteria by 14-dpi (63). In contrast, however, the *S. aureus* BIAI strain 117 displayed complete recalcitrance to TAPI, with high CFUs from harvested implants or tissues and no significant difference in CFUs recovered from samples of TAPI treated compared to control mice over a 14-d time course. Together, these data suggest that TAPI is only effective at reducing CFUs at early time points after surgery and that any bacteria that persist in the presence of TAPI can go on to cause chronic infection. Additionally, these data suggest that while TAPI may have some efficacy against *P. aeruginosa* strains causing BIAI, as BIAI strain 157 was fairly susceptible to TAPI in the mouse model, the irrigant may not be as efficacious at preventing BIAI with *S. aureus*, as the *S. aureus* isolates, and particularly the BIAI strain 117, were uniquely persistent over a 14-d time course. Most importantly, these data highlight the discrepancies between *in vitro* and *in vivo* results, and suggest they may not always accurately inform the efficacy of prevention strategies.

In seeking to understand why the *S. aureus* strains displayed increased recalcitrance *in vivo* compared to the *P. aeruginosa* isolates, despite exhibiting increased susceptibility to the antibiotics in TAPI, we assessed the ability of these strains to form community structures. The most well studied bacterial communities are biofilms, which promote recalcitrance to antibiotics as well as the host immune system (35, 36, 43, 44). All *S. aureus* and *P. aeruginosa* isolates were able to form biofilm under *in vitro* conditions. Importantly, these biofilms provided increased recalcitrance to antibiotic concentrations at which the planktonic bacteria were susceptible. Fortunately, TAPI was effective at significantly reducing the biomass of the biofilm formed by all three *P. aeruginosa* strains. In contrast, *S. aureus* biofilm and TAPI recalcitrance was affected by the conditions used to form biofilm. Specifically, using an *in vitro* biofilm assay that more closely mimics the host environment during infection, we demonstrated that when *S. aureus* formed biofilm in the presence of human plasma, the biomass increased along with the pathogen’s recalcitrance to TAPI. *S. aureus* is known to exploit host proteins released as part as the inflammatory and wound healing pathway, such as fibrinogen and collagen, and incorporate them into their biofilm structure to promote recalcitrance (36, 64-66). These host proteins, which accumulate within the breast during the surgical procedures required for implantation of breast prostheses and attach to the device surface, create a suitable environment for *S. aureus* biofilm formation during BIAI (24, 25, 36). Furthermore, it is well known that antibiotics themselves can act as signals to induce biofilm formation or promote drug tolerance (67-69). Thus, these data suggest that *S. aureus* isolates causing BIAI may encode unique mechanisms that respond to host proteins or antibiotic stimuli to promote recalcitrance during infection compared to reference strains, which are commonly used for these types of studies.

This study demonstrates that reference strains can exhibit fundamentally different phenotypes compared to clinically relevant, currently circulating BIAI isolates. Specifically, while the *S. aureus* BIAI strain 117 displayed similar MICs to the reference strain JE2, the BIAI isolate displayed increased recalcitrance to TAPI during biofilm formation and *in vivo*. Furthermore, the discovery that TAPI may increase the biofilm biomass of the *S. aureus* BIAI isolates has important implications for prophylactic antibiotic treatment in breast implant-based reconstructive surgeries, as TAPI may provide the signals to promote BIAI with this pathogen. Overall, these results emphasize the need for additional studies with more clinically relevant strains to fully understand the mechanisms these pathogens encode to cause chronic BIAI. Additionally, there should be a sense of urgency in updating current protocols to include better antibiotic stewardship practices to prevent the overuse and misuse of antibiotics. Finally, future studies that dissect the pathogenic mechanisms that promote recalcitrance among BIAI are sorely needed to develop targeted therapies that can either effectively prevent or eradicate these chronic infections.

## Materials and Methods

### Strains and Growth Conditions

All strains used in this study are listed in **Table 1** (51, 70). The *S. aureus* and *P. aeruginosa* isolates that caused BIAIs were provided by Dr. Margaret Olsen at Washington University in St. Louis School of Medicine. Brain heart infusion (BHI) broth (BD, #237200) and agar (BD, #214010) plates were used to maintain and prepare all cultures for experiments. For the MIC and MBC assays, bacterial isolates were grown in Mueller Hinton broth (MHB) (Sigma; #7019-100G). For animal experiments, cells were harvested from overnight cultures grown at 37°C, shaking in BHI broth.

### MIC and MBC Assays

MIC and MBC assays were performed following the Clinical Laboratory Standards Institute (CSLI) guidelines and as we have previously published (24, 71, 72). Briefly, *S. aureus* was grown to an OD_600_ of 0.4, which corresponds to ∼2.8×10^8^ CFU/mL, and then diluted to ∼1×10^6^ CFU/mL. *P. aeruginosa* was grown for 4 hours and diluted once to achieve ∼1×10^6^ CFU/mL. Individual antibiotics, including bacitracin (ThemoFisher, #226100050), cefazolin (ThermoFisher, #455210010), and gentamicin (ThermoFisher, #455310050), were diluted to concentrations ranging from 0.0625 to 2800 ug/mL in MBH. Antibiotics were then added to each bacterial suspension at a 1:1 ratio (100 uL bacteria:100 uL antibiotic) in a 96-well plate (Fisher Scientific, #07-000-108) resulting in a final concentration of 5×10^5^ CFU/mL of bacteria and antibiotics ranging from 0.03125-1400 ug/mL. Controls included MHB medium alone, bacteria in MHB medium alone, and MHB with antibiotics alone. The 96-well plates were then incubated overnight at 37°C. The OD_600_ was measured using a Synergy H1 Biotek microtiter plate reader and the MICs were determined based on an OD_600_ value of less than 0.1. CFUs were enumerated from the MIC assays to determine the MBCs, which are based on a 99% reduction of CFUs. Each experiment contained three and was repeated at least twice. MIC breakpoints were obtained from CLSI for gentamicin and cefazolin, and knowledgebase for bacitracin and are as followed for *S. aureus:* ≥1 ug/mL for gentamicin, ≥4 ug/mL for cefazolin, and ≥ 8 ug/mL for bacitracin and for *P. aeruginosa*: ≥16 ug/mL for gentamicin, ≥ 8 ug/mL for cefazolin, and 700 ug/mL for bacitracin (71, 73). Additionally, all three antibiotics were combined into the TAPI solution used in patients: 80 mg gentamicin, 1 g Cefazolin, and 50,000 U bacitracin in 500 ml saline (50). Susceptibility to the TAPI solution was measured using the same procedure described above for the MIC and MBC assays. Each experiment contained three replicates and was repeated at least twice.

### WGS of the S. aureus and P. aeruginosa BIAI Isolates

Short and long read sequencing methods were used to WGS all BIAI isolates as previously described (74, 75). Genomes are available under NCBI BioProject # PRJNA859168. Briefly, the *S. aureus* and *P. aeruginosa* strains were grown in BHI and LB, respectively, for 3 hours, shaking at 37°C. The cultures were spun down at 10,000 g for 5 mins, the supernatant discarded, and the genomic DNA was then extracted using the Qiagen QIAamp DNA Mini Kit (Qiagen, cat # 51306) following the manufacturer’s protocol, with the exception that 1 ug/ml of lysostaphin and 20 ug/mL of lysozyme was added to the initial buffer. Sequencing was performed as previously described (76). Briefly, genomic DNA underwent library preparation, using the SQK-RBK004 library preparation kit, and long-read sequencing via the Oxford Nanopore GridION X5 (Oxford, UK) sequencer. Additionally, short read sequencing was performed using the Illumina NextSeq 2000 sequencer. Bacterial genomes were then assembled as previously described (76). Briefly, raw assemblies were created with Flye-v2.7, and then underwent error correction and polishing steps using Racon-v-1.4.5 and Medaka-v0.11.5. Completed genomes were analyzed for antimicrobial gene carriage using Abricate (https://github.com/tseemann/abricate) and the CARD resistance database (77).

### Biofilm Assays

Biofilm assays for *S. aureus* and *P. aeruginosa* were performed as previously described (29, 35). Briefly, an overnight culture of each strain, grown in BHI for *S. aureus* or LB for *P. aeruginosa*, was subcultured to an OD_600_ of 0.2 and then further diluted 1:100 in fresh BHI or LB, respectively. Next, 200 uL of the subculture was added to a 96-well plate and incubated, shaking at 37°C overnight to allow biofilm to form. The next day the supernatant culture was then carefully removed from the top of the biofilms and 200 uL of the antibiotic solution, which was prepared as described above and used at the concentration previously determined as the MIC, twice the MIC, and four times the MIC for each strain, was added. The plate was then incubated, shaking at 37°C for 18-20 hours. The next day the supernatant culture was removed and the remaining biofilm was air dried for 30 minutes, washed with sterile water, stained with 0.1% crystal violet, washed again to remove excess crystal violet, and then the remaining crystal violet was solubilized with 33% acetic acid. The OD_600_ was then measured using a Synergy H1 Biotek microtiter plate reader. Additionally, *S. aureus* biofilm was also formed in the presence of 20% human plasma. For these experiments human plasma (Sigma; #P9523-5ML) was diluted to 20% in NaHCO3 (Sigma; #S8875) and added to a 96-well plate, which was then incubated at 4°C overnight, as previously described (29). NaHCO3 coated wells were used as a negative control. The next day the supernate was removed and the *S. aureus* biofilm assay was performed as described above.

### Aggregation Assays

The *P. aeruginosa* aggregation assay was performed as previously published (35), with some modifications. Briefly, overnight cultures of *P. aeruginosa* grown in LB were diluted 1:100 in fresh LB. Aggregates were then formed by growing the strains statically at 37°C for 24 hrs. A 1 uL sterile loop (BD, cat# 220215) was used to collect uniform amounts of aggregates, which were carefully suspended in 90 uL of PBS in a 96-well plate. Gentamicin was prepared as described above and added to the wells. Aggregates without antibiotics were used as controls. The plate was then incubated statically, overnight at 37°C. The next day, the plate was centrifuged for 8 minutes at 3500 rpm to sediment the aggregates in the wells. The supernate was removed and all remaining aggregates were processed, stained with crystal violet, and the OD measured as described above.

### Mouse model of BIAI

One representative BIAI isolate (117 and 157) and one reference strain (JE2 and PAO1) for each bacterial species was selected for these studies. The mouse model was performed as previously described (31, 70). Briefly, ∼10^7^ CFUs of *S. aureus* and ∼10^5^ CFUs of *P. aeruginosa* were prepared in 1X phosphate buffered saline (PBS). Mice were anesthetized via 1-4% isoflurane inhalation, the hair was removed from an area of the back, and the site of surgery was disinfected using chlorohexidine and 70% isopropyl alcohol. After creating a small (∼1 cm) subcutaneous pocket in the haunch of the mouse, 100 uL of either sterile saline (McKesson, cat# 186661) or the TAPI (prepared as described above) was administered directly into the pocket via a 1 mL syringe (BD, cat# 309628). The implants, which were prepared by using a 6 mm Integra Miltex Standard punch to cut small, standardized pieces from the shell of a NATRELLE INSPIRA® SoftTouch (Ref# SSM-755) smooth silicone breast implant, were UV sterilized for at least 1-hour before they were inserted into the subcutaneous pocket. The incision was then closed using sutures or wound clips. Each pocket was infected with the prepared bacterial inoculum via subcutaneous injection in the area where the implant was visible under the skin. Implants and tissue samples near the implants were harvested at 1-, 7-, and 14-dpi and CFUs were determined as previously described (29, 31, 70). Briefly, the implants were sonicated in 1X PBS for 10 minutes and vortexed for a 1 minute, twice before serial dilution and plating on agar plates. Additionally, tissue samples were weighed and then homogenized using a MP-BIO FastPrep24 bead beater for 1 minute, twice with a 5-minute rest in between before serial dilution and plating, as described above. All animal work was approved by the Animal Welfare Committee (protocol # AWC-20-0057).

### Statistical analyses and sample size calculations

The Mann Whitney U test was used to determine significant differences biomass recovered between community formation treated with antibiotics compared to no antibiotic controls as well as to determine significant differences between the CFUs recovered from mice that received TAPI compared to those that received saline. All statistical tests were performed using GraphPad Prism 8.4.3 software.

## Acknowledgments

We would like to thank Dr. Margaret Olsen at Washington University in St. Louis School of Medicine for providing the BIAI isolates, Samantha Hitt for making the implant punches used in this study, and An Dinh and Shelby Simar for their sequencing advice. Dr Myckatyn receives royalties for product development, funds for an investigator initiated trial associated with acellular dermal matrices in breast reconstruction, and advisory board remuneration from RTI Surgical. He receives an investigator initiated award from Sientra that studies the metabolomics of breast tissue expander infection. None relate directly to the topic matter of this study and no industry funds were received for completing this study. He has no other disclosures. All other authors declare no other conflicts of interest.

